# GenPept-Curated-2025: A Benchmark Dataset for Antimicrobial Peptide Prediction with Homology-Controlled Partitioning

**DOI:** 10.64898/2026.04.25.720793

**Authors:** Huynh Trong Pham, Bao Huynh, Thanh-Hoang Nguyen-Vo

## Abstract

Antimicrobial peptides (AMPs) are promising therapeutic candidates against rising antimicrobial resistance, yet progress in AMP prediction is hampered by the lack of benchmark datasets that address homology leakage, negative set reliability, and distributional diversity. Existing AMP databases, designed as biological repositories, do not enforce the controlled partitioning required for rigorous machine learning evaluation. We present `GenPept-Curated-2025`, a curated, class-balanced benchmark of 11,000 peptide sequences (5,500 AMP / 5,500 non-AMP) derived from Bacteria, Archaea, and Fungi, and sourced exclusively from GenPept/NCBI Protein. The dataset was constructed through a reproducible pipeline comprising taxonomic scoping, quality control, precursor handling, annotation-based labeling, and Identical Protein Groups (IPG)-based deduplication, with sequence length restricted to 10–200 aa. The AMP proportion varies substantially across length bins (14.2% in [10, 50] aa to 77.1% in [101, 150] aa), identifying length-dependent class imbalance as a distribution shift that benchmarking must account for. The dataset is openly released to support standardized, reproducible, and leakage-free evaluation of AMP prediction models.

## 1 Introduction

### 1.1 Background

Bacterial antimicrobial resistance has become one of the most pressing global health challenges. In 2021, approximately 4.71 million deaths were associated with, and 1.14 million were directly attributable to, multidrug-resistant bacterial infections, with the burden concentrated in low- and middle-income countries.^1^ Concurrently, the antimicrobial development pipeline shows signs of stagnation: the WHO’s 2025 pipeline report recorded a decline from 97 agents in clinical development in 2023 to 90 as of February 2025, of which only 15 are considered truly innovative and only five are effective against pathogens in the updated 2024 “critical” priority list. Since July 2017, only 17 agents targeting priority pathogens have been approved, with just two representing entirely new chemical classes. This dual crisis of insufficient quantity and limited innovation underscores the urgent need for non-traditional therapeutic strategies, among which antimicrobial peptides have emerged as a highly promising approach.

### 1.2 Limitations of Existing Data Resources

Artificial intelligence and deep learning have substantially advanced AMP prediction, from protein language models^2^ to generative systems capable of discovering novel bioactive molecules. ^3^ Yet model performance and generalizability remain fundamentally constrained by dataset quality.^4^

The AMP database ecosystem has expanded rapidly, with APD6, DBAASP v3, ^5^ CAMPR4,^6^ DRAMP 4.0,^7^ and dbAMP 3.0^8^ collectively providing curated data on peptide activity, toxicity, structure, and biochemical properties.^9^ However, these resources are designed as biological repositories rather than machine learning benchmarks: they do not enforce homology-controlled partitioning, provide experimentally validated negative sets, or ensure reproducible train/test splits across diverse sequence distributions.

### 1.3 Motivation and Objectives

The limitations outlined above collectively point to an unmet need: a purpose-built AMP benchmark that addresses homology leakage, negative set quality, and distributional diversity within a unified, reproducible framework. `GenPept-Curated-2025` is designed to fill this gap via a systematic extraction–cleaning–normalization pipeline applied to GenPept, homology-aware cluster-based partitioning, and carefully curated negative set construction, enabling rigorous, leakage-free evaluation of AMP prediction models across diverse sequence distributions. The contributions of this work are as follows:

- **Comprehensive curation:** A reproducible extraction–cleaning–normalization pipeline sourced from GenPept, ensuring sequence quality, annotation consistency, and label reliability across positive and negative sets.
- **Homology-controlled evaluation:** A cluster-based partitioning protocol that controls sequence similarity across splits, mitigating data leakage and enabling evaluation outcomes that more faithfully reflect real-world generalizability.
- **Diversity-aware benchmarking:** Diverse sequence distributions across varying lengths and biological origins, supporting realistic assessment of model robustness in AMP discovery scenarios.
- **Community utility:** `GenPept-Curated-2025` is openly released as a benchmark resource supporting standardized, comparable, and reproducible evaluation across future AMP prediction studies.

## 2 Materials and Methods

### 2.1 Overview of the Curation Workflow

`GenPept-Curated-2025` was constructed through a reproducible multistage curation workflow designed to improve source consistency, reduce redundancy, and increase label reliability for AMP benchmarking. Briefly, protein records were retrieved from GenPept within a pre-defined taxonomic scope (*Bacteria, Archaea*, and *Fungi*), followed by sequence-level quality control enforcing the target length range, removal of non-standard amino acids, and exclusion of unreliably annotated entries. Precursor records were flagged separately to distinguish uncertain cases from the main benchmark set. Annotation-based labeling was applied to the DEFINITION field and, where available, the linked CDS product field to assign AMP and non-AMP labels while filtering known false-positive naming patterns. Redundancy was resolved using Identical Protein Groups (IPG), ensuring each sequence was represented only once. The curated dataset was then summarized across length intervals to characterize class composition and associated uncertainty. A schematic summary of the workflow is provided in Figure 1.

**Figure 1:**
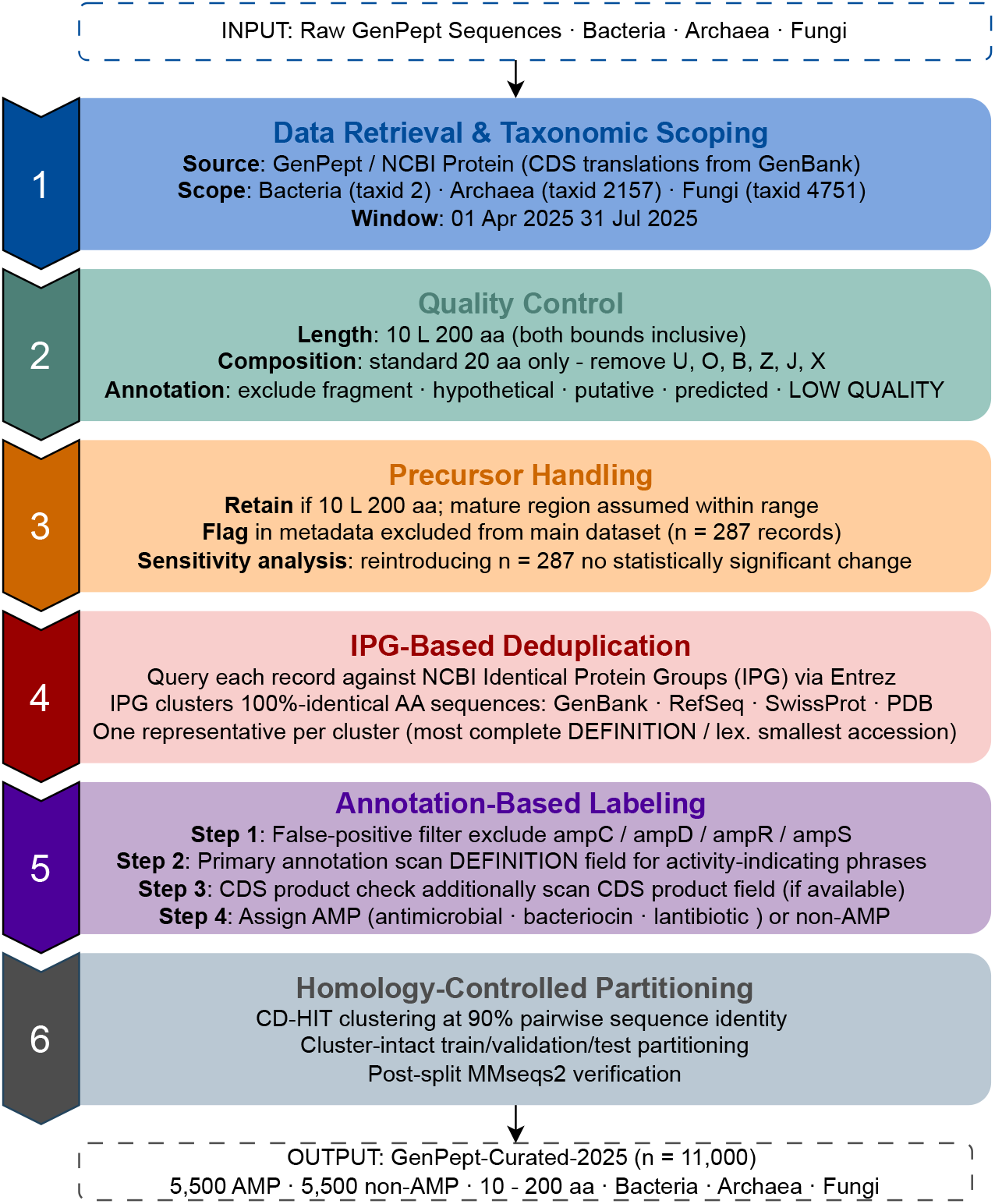
Schematic overview of the GenPept-Curated-2025 curation pipeline.

### 2.2 Data Retrieval and Taxonomic Scoping

#### 2.2.1 Data Sources

Sequences were retrieved from the NCBI Protein database via GenPept, which collects translations of coding sequence (CDS) regions deposited in GenBank.^10^ RefSeq Protein was used solely for identity cross-referencing and contributed no sequences to the benchmark, ^11^ preserving source consistency and reducing cross-database redundancy.

#### 2.2.2 Taxonomic Scope

The dataset is restricted to Bacteria (taxid 2), Archaea (taxid 2157), and Fungi (taxid 4751), focusing on microbe-derived AMPs of high relevance to antimicrobial resistance (AMR) while reducing distributional heterogeneity relative to viral, animal, or plant peptides. Viruses (taxid 10239), Metazoa (taxid 33208), Embryophyta (taxid 3193), and other non-fungal Eukaryota are excluded. All sequences were retrieved over the window 01/04/2025–31/07/2025.

#### 2.2.3 Operational Definition and Length Criteria

The term *peptide* is operationally defined as any polypeptide satisfying 10 *≤ L ≤* 200 amino acids (inclusive), independent of biological activity. Sequences are labelled **AMP** if they carry recorded antimicrobial activity, or **non-AMP** if no such record exists; the non-AMP label reflects *absence of evidence*, not *evidence of absence*. The upper bound of 200 aa exceeds thresholds used by some databases (e.g., DRAMP 4.0 and dbAMP 3.0 apply *<*100 aa), and is justified on three grounds:

i. To cover microbial propeptide and precursor sequences whose mature regions are often *<*100 aa but whose cleavage boundaries may be incompletely annotated.
ii. To maintain consistency with the preprocessing pipeline, where the padded input length is fixed at *L*_pad_ = 200, avoiding separate workflows for peptides and small proteins.
iii. To preserve the full length distribution and reduce selection bias during dataset construction.

The lower bound *L ≥* 10 excludes oligopeptides rarely reported to exhibit independent antimicrobial activity. Together, the 10–200 aa range aligns with existing AMP database practices and provides a uniform input space for model comparison.

### 2.3 Quality Control

Filtering criteria follow QC practices reported in recently updated AMP databases.^7–9^ Only sequences satisfying all three conditions below were retained.

#### Condition 1

**Length**. 10 *≤ L ≤* 200 aa, both bounds inclusive.

#### Condition 2

**Amino acid**. Only the 20 standard amino acids (ACDEFGHIKLMNPQRSTVWY) are permitted.^12^ Sequences containing non-standard characters (`U, O, B, Z, J, X`), residual hyphens, or other artifacts are removed during GenPept translation field parsing, with whitespace normalization applied in the same step.

#### Condition 3

**Annotation quality**. Records with unreliable `DEFINITION` field descriptions are excluded, with the linked CDS `product` field additionally checked where available. Exclusion targets the keywords *fragment, hypothetical, putative, possible, predicted*, and *LOW QUALITY PROTEIN*, matched case-insensitively as full phrases to prevent spurious substring matches.

This QC step reduces indirect label noise and limits annotation-driven bias in model evaluation.

### 2.4 Precursor Handling

Records annotated as precursor sequences without mature-peptide boundary information are retained if 10 *≤ L ≤* 200, on the assumption that the potential mature segment falls within the defined range. These records are flagged in the metadata and excluded from the main dataset (*n* = 11,000) to avoid label noise. A sensitivity analysis reintroducing 287 flagged precursor records showed no statistically significant differences in model performance or conclusions, confirming the robustness of this approach.

### 2.5 IPG-Based Deduplication

Each GenPept record is identified by `accession.version` and queried against NCBI IPG via Entrez. `IPG` clusters protein records with 100% identical amino-acid sequences from different sources (GenBank, RefSeq, SwissProt, PDB, etc.) under a shared `IPG_ID`. Within each group, a single representative is retained by: (i) preferring the record with the most complete `DEFINITION` field; and (ii) selecting the lexicographically smallest `accession.version` when completeness is comparable. Deduplication is performed before labeling and before CD-HIT clustering, ensuring each unique sequence appears exactly once.

For prokaryotes (Bacteria and Archaea), the `WP_` non-redundant protein accession is recorded when available in IPG.^11^ For Fungi, where `WP_` does not apply, `NP_` or `XP_` accessions are recorded when present, or the RefSeq field is left blank. All cross-reference information is retained in the metadata.

### 2.6 Annotation-Based Labeling

Labeling follows curated-annotation practices used in major AMP databases, including APD6,^9^ dbAMP 3.0,^8^ DRAMP 4.0^,7^ and CAMPR4^,6^ summarized in four steps:

#### Step 1. False-positive filtering

Records whose gene or protein name contains `amp` but is unrelated to antimicrobial activity—specifically `ampC, ampD, ampR`, and `ampS`—are removed.

#### Step 2. Primary annotation check

A record is identified as AMP if the `DEFINITION` field contains activity-indicating phrases.

#### Step 3. CDS product check

Where a CDS link exists, the product field is additionally checked under the same rules.

#### Step 4. Label assignment

Each sequence is assigned **AMP** or **non-AMP** based on Steps 1–3.

The pattern-matching lexicon covers two groups: (i) activity terms—*antimicrobial peptide, antibacterial peptide, antifungal peptide, antiparasitic peptide*, accepting hyphenated variants; and (ii) microbial peptide classes—*bacteriocin, lantibiotic, microcin, thiopeptide, sactipeptide*.^7,9^ All matching is case-insensitive with word-boundary constraints to prevent false positives from unrelated substrings (e.g., *amphiphysin*). Residual label noise was further reduced by the gene-name filter in Step 1 and, when needed, by manual inspection of randomly sampled ambiguous records.

### 2.7 Homology-Controlled Partitioning

Data leakage from high sequence similarity across data partitions can inflate performance estimates and obscure true generalization. To mitigate this, benchmark partitions were generated through a three-stage procedure: clustering, cluster-level partitioning, and post hoc verification. Throughout this study, homology-controlled denotes cluster-aware control of high-identity cross-split redundancy under an explicit 90% identity criterion.

- **Stage 1. Clustering.** Peptide sequences were clustered using CD-HIT 4.8.1 at 90% pairwise sequence identity, with a coverage threshold of 0.80 on the shorter sequence.
- **Stage 2. Cluster-level partitioning**. Entire clusters were assigned to a single partition with training (n = 7,700), validation (n = 990), or test (n = 2,310) sets, corresponding to a 70:9:21 split ratio. Class balance across AMP/non-AMP labels was maintained, and a fixed random seed (42) was used for reproducibility.
- **Stage 3. Post hoc verification**. Post hoc verification. An all-vs-all MMseqs2 search across partition pairs confirmed the absence of cross-split leakage: no sequence pair met both thresholds of *≥*90% identity and *≥*80% coverage of the shorter sequence.

## 3 Results and Discussion

After applying all curation steps, the main dataset contains *n* = 11,000 sequences (5,500 AMP and 5,500 non-AMP) (Table 1). Table 2 reports label-wise counts and AMP proportions across four length intervals, with uncertainty quantified by the Wilson score interval,^13^ which provides more stable bounds than the Wald approximation under unequal subgroup sizes and proportions deviating from 0.5^14^—both conditions present here. Figure 2 visualizes the interval-specific AMP proportions with their 95% Wilson confidence intervals (CI).

**Table 1:**
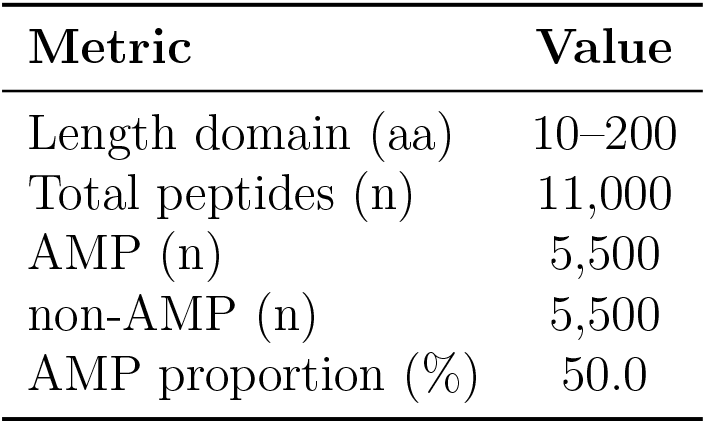
Dataset size after the data preparation stage.

**Table 2:** Distribution of peptide counts and AMP proportion by length interval in GenPept-Curated-2025.

| Length interval (aa) | Total | AMP | non-AMP | AMP proportion (%) | Wilson 95% CI |
| --- | --- | --- | --- | --- | --- |
| 10–50 | 2,618 | 372 | 2,246 | 14.2 | [12.9–15.6] |
| 51–100 | 2,379 | 1,602 | 777 | 67.3 | [65.4–69.2] |
| 101–150 | 2,281 | 1,759 | 522 | 77.1 | [75.3–78.8] |
| 151–200 | 3,722 | 1,767 | 1,955 | 47.5 | [45.9–49.1] |
| **Total** | **11,000** | **5,500** | **5,500** | **50.0** | **[49.1–50.9]** |

*Note:* Intervals use closed boundaries. The AMP proportion is computed as AMP / (AMP + non-AMP) within each interval. The 95% CI is estimated using the Wilson method.

**Table 3:** Files included in the GenPept-Curated-2025 repository.

| File | Description |
| --- | --- |
| `dataset.csv` | 11,000 sequences with full metadata |
| `amp_sequences.fasta` | 5,500 AMP sequences, FASTA format |
| `nonamp_sequences.fasta` | 5,500 non-AMP sequences, FASTA format |
| `train.csv` | Training split (CD-HIT 90% identity) |
| `test.csv` | Test split (CD-HIT 90% identity) |

**Figure 2:**
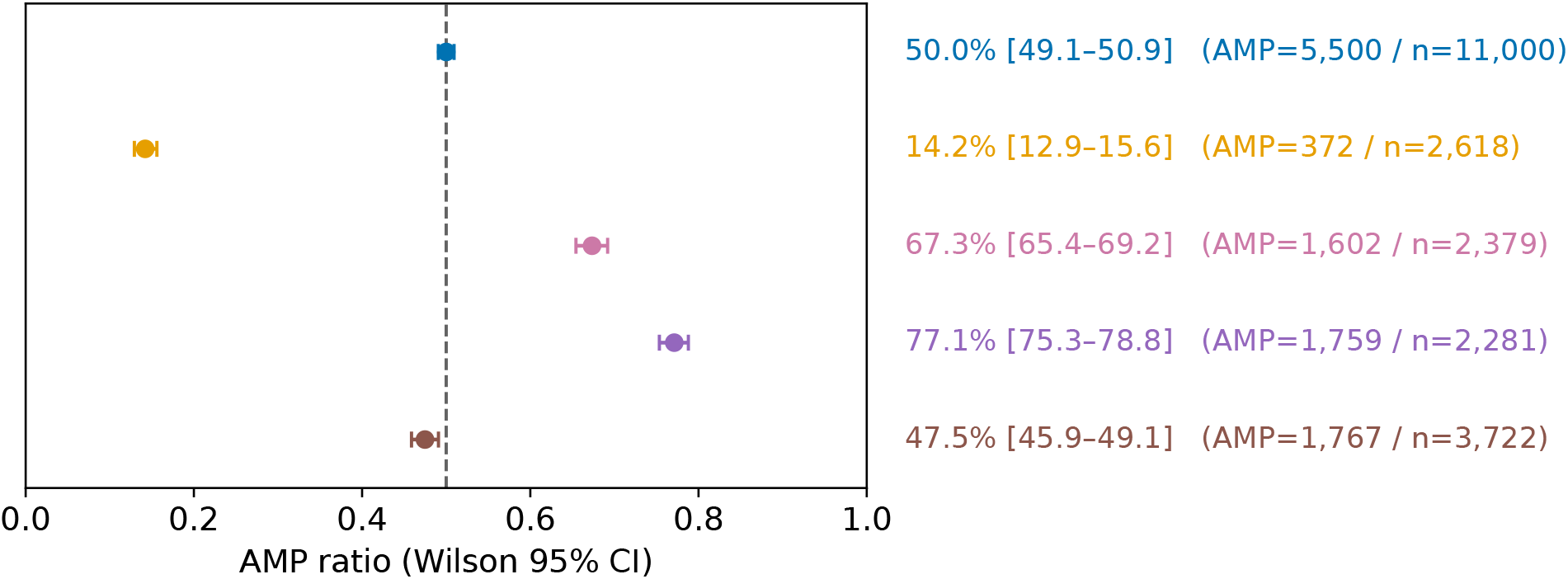
AMP proportion by length interval with 95% Wilson confidence intervals. Each point represents one length bin; the dashed line marks the global AMP proportion (50%). Sample sizes and AMP counts are annotated per bin.

The AMP proportion varies substantially across length bins, from 14.2% in [10, 50] aa to 77.1% in [101, 150] aa. Non-overlapping Wilson CIs across most bin pairs confirm these differences exceed sampling variability. The [51, 100] aa and [101, 150] aa intervals are markedly enriched for AMP sequences, while [10, 50] aa is dominated by non-AMP sequences. Length-dependent class imbalance is therefore a structural property of the dataset that must be accounted for in benchmark interpretation.

The AMP proportion varies substantially across length bins (14.2% in [10, 50] aa vs. 77.1% in [101, 150] aa), confirming a non-uniform class distribution that constitutes a source of distribution shift requiring control in benchmarking. Figure 3 illustrates this length-dependent class structure: Panel A shows that absolute AMP and non-AMP counts vary markedly across intervals, while Panel B normalizes by interval size to isolate class composition from sample-size effects, revealing that the [10, 50] aa interval is non-AMP-dominated, the [51, 150] aa range is AMP-enriched, and the [151, 200] aa interval shows a more balanced composition.

**Figure 3:**
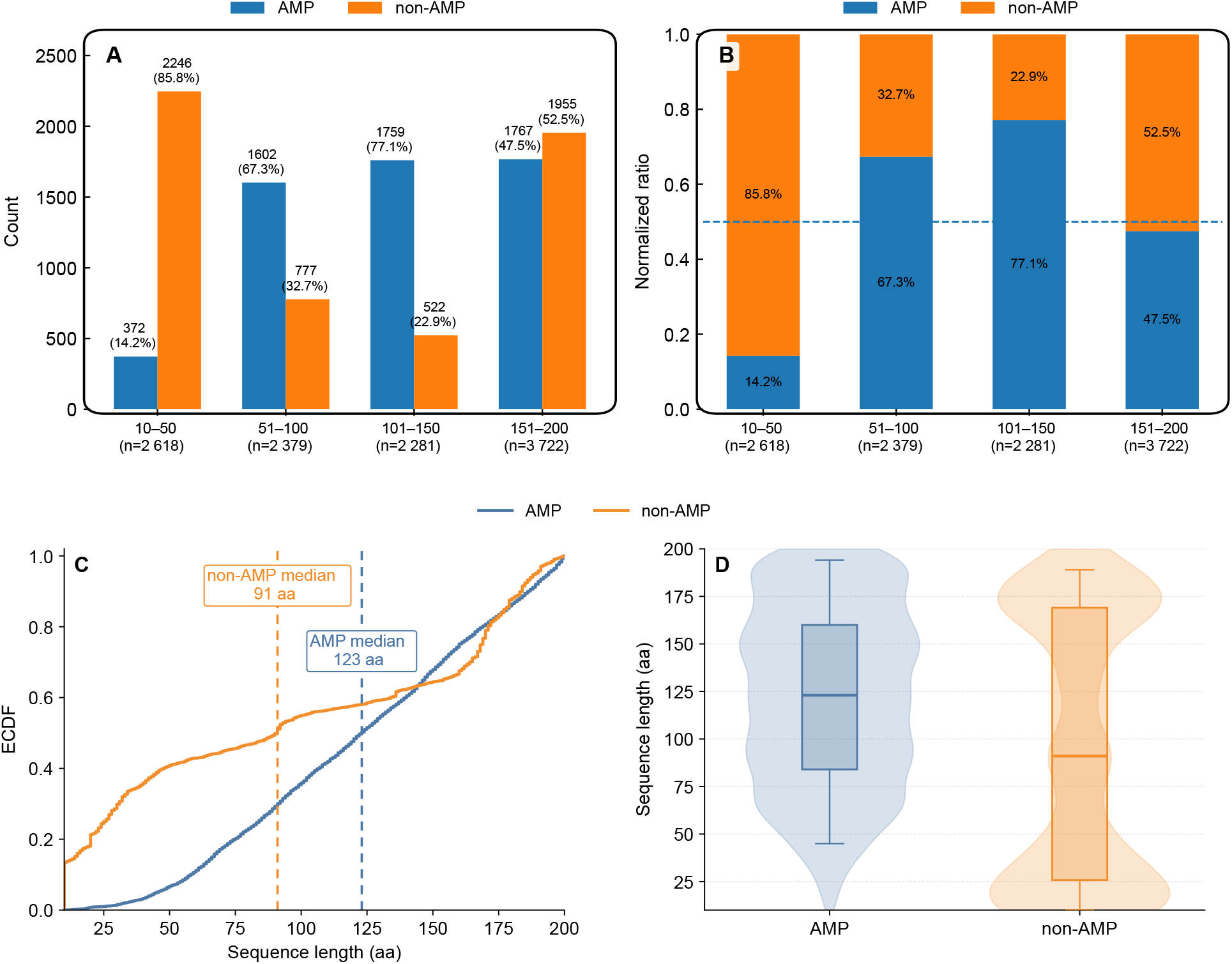
Class composition across the length domain. (A) AMP proportion per length interval with 95% Wilson confidence intervals; each point represents one length bin, sample sizes and AMP counts are annotated per bin, and the dashed line marks the global AMP proportion (50%). (B) Absolute AMP and non-AMP counts per interval. (C) Normalized proportions; the dashed line marks the global AMP proportion (50%).

This pattern is consistent with the interval-specific proportions reported in Table 2. Together, these observations indicate that class imbalance is length-dependent rather than globally constant, and benchmark outcomes should therefore be interpreted in the context of the underlying length distribution.

Figure 3 extends the interval-level summaries to full empirical length distributions for each class. The ECDF curves show that AMP and non-AMP sequences diverge across a broad span of the length domain rather than at a single cutoff, while the violin-boxplot highlights differences in central tendency and spread. Together, these patterns indicate that sequence length is a structural characteristic of `GenPept-Curated-2025` that may systematically affect model behavior and should therefore be considered when interpreting evaluation results.

## 4 User Guide

The main benchmark table contains curated AMP and non-AMP sequences, each identified by accession number, amino acid sequence, class label, and taxonomic origin. Supplementary metadata files report processing flags assigned during curation, including precursor status and cross-reference information where available, documenting the filtering, labeling, and deduplication decisions underlying each record.

### 4.1 Accessing the Dataset

`GenPept-Curated-2025` is publicly available at: https://github.com/biochem-data-sci/GenPept-Curated-2025. The repository contains the following files:

Key columns in `dataset.csv`: *accession* (GenPept accession.version), *sequence* (amino acid string), *label* (AMP / non-AMP), *length* (integer, 10–200), *taxon* (Bacteria / Archaea / Fungi), *ipg_id* (NCBI IPG cluster ID), *refseq_accession* (`WP_` / `NP_` / `XP_`when available).

### 4.2 Loading in Python Environment

The `train.csv` and `test.csv` splits are controlled at 90% sequence identity via CD-HIT; resplitting randomly will introduce homology leakage and should be avoided. The non-AMP label reflects absence of annotation evidence rather than confirmed biological inactivity. Length-dependent class imbalance should be accounted for during training, as the AMP proportion ranges from 14.2% ([10, 50] aa) to 77.1% ([101, 150] aa), making stratified sampling or length-aware evaluation advisable. The `precursor_flagged.csv` file is reserved for sensitivity analysis and should not be included in standard training. The dataset can be loaded in Python with the script shown in Figure 4. Users are asked to cite this paper when using `GenPept-Curated-2025` in published work.

**Figure 4:**
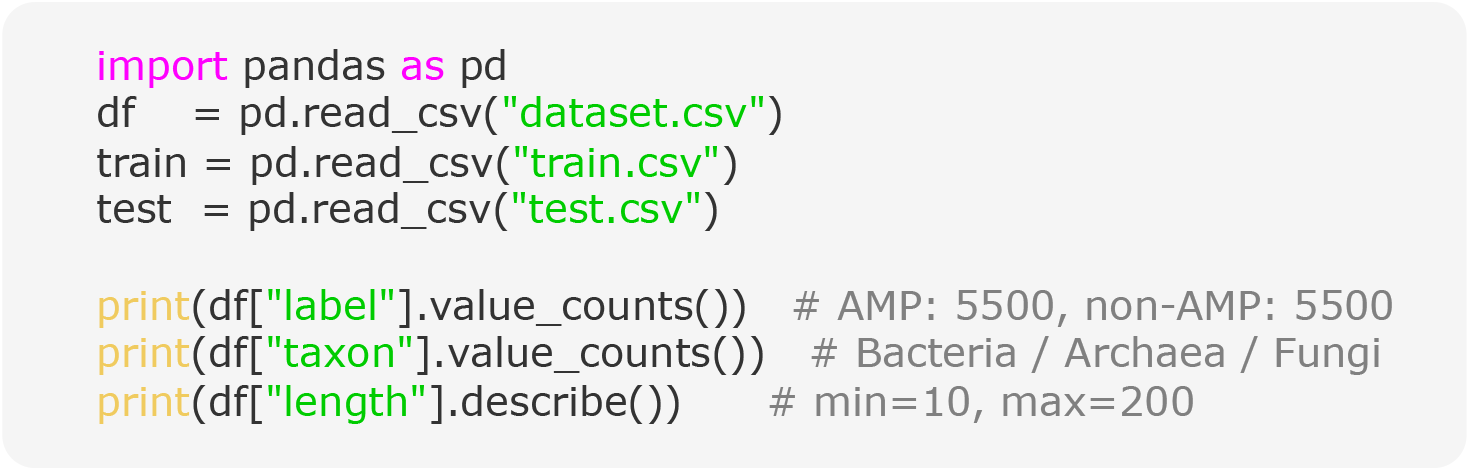
Loading datasets in Python environment.

## 5 Conclusion

We presented `GenPept-Curated-2025`, a curated benchmark of 11,000 class-balanced sequences (5,500 AMP / 5,500 non-AMP) from *Bacteria, Archaea*, and *Fungi*, designed to address homology leakage, unreliable negative set construction, and inadequate distributional reporting. The pipeline applies IPG-based deduplication, annotation-quality filtering, and precursor flagging to reduce label noise and cross-record redundancy reproducibly. Characterization reveals substantial length-dependent variation in AMP proportion, confirming that length distribution is a structural property that can bias benchmark outcomes if uncontrolled.The dataset complements existing AMP databases by making benchmarking assumptions explicit, with the non-AMP label interpreted under the “absence of evidence” principle. We anticipate that incorporating experimentally validated negatives and broader taxonomic coverage will further strengthen this benchmarking framework.

## Author’s contributions

**H.T.P**.: Data Curation, Methodology, Investigation, Formal analysis, Writing - Original Draft. **B.H**.: Conceptualization, Formal analysis, Writing - Review & Editing, Supervision. **T.-H.N.-V**.: Conceptualization, Formal analysis, Writing - Review & Editing, Supervision.

## Note

The authors declare no competing financial interest.

## Data and Software Availability

The link to the code with data used in this study are available at https://github.com/biochem-data-sci/GenPept-Curated-2025.

## TOC Graphics

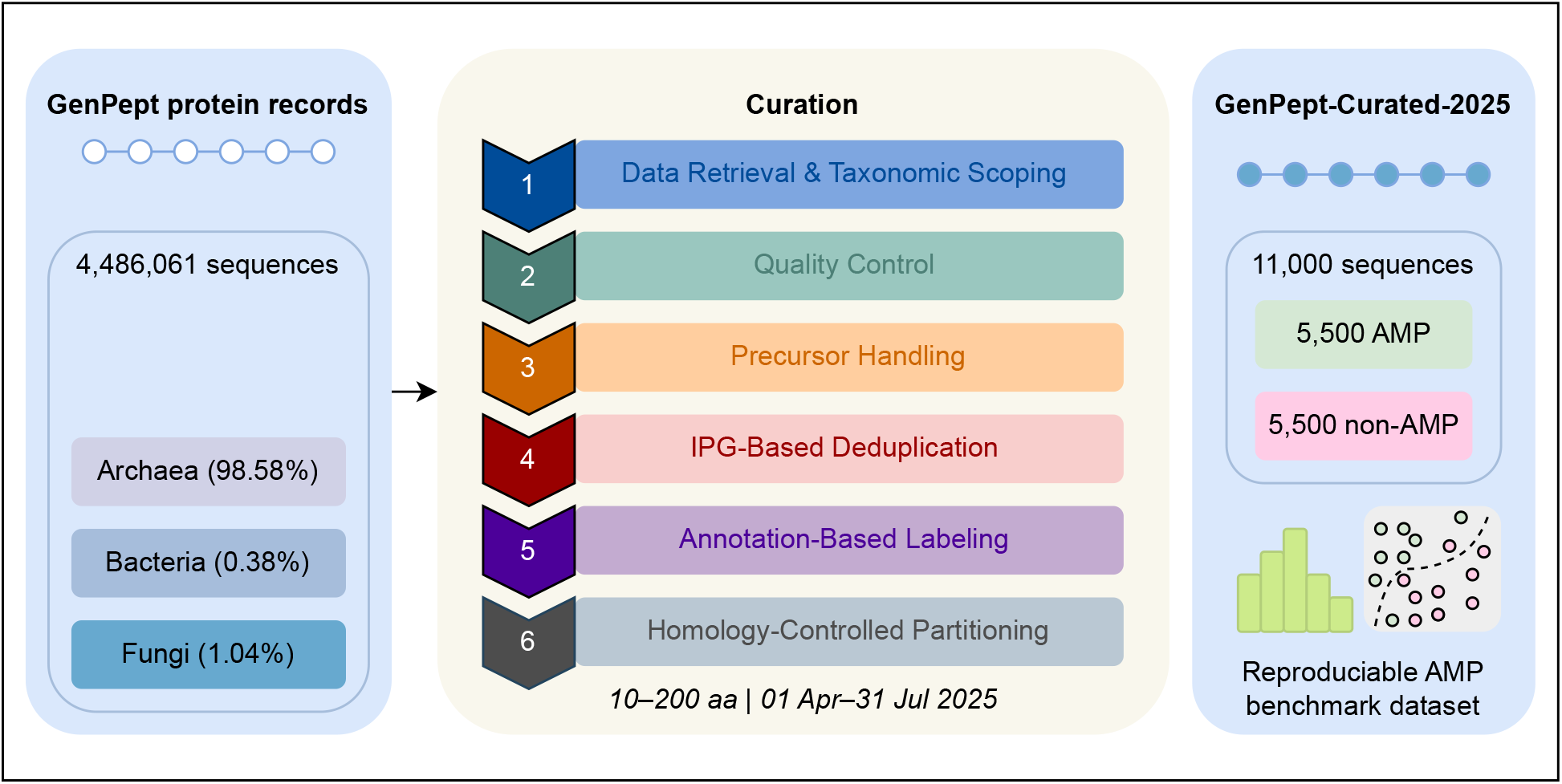

